## Supplementary material for "Surveillance on California dairy farms reveals multiple possible sources of H5N1 transmission": Figures S1, S2 and S3

### Supporting Information Figures

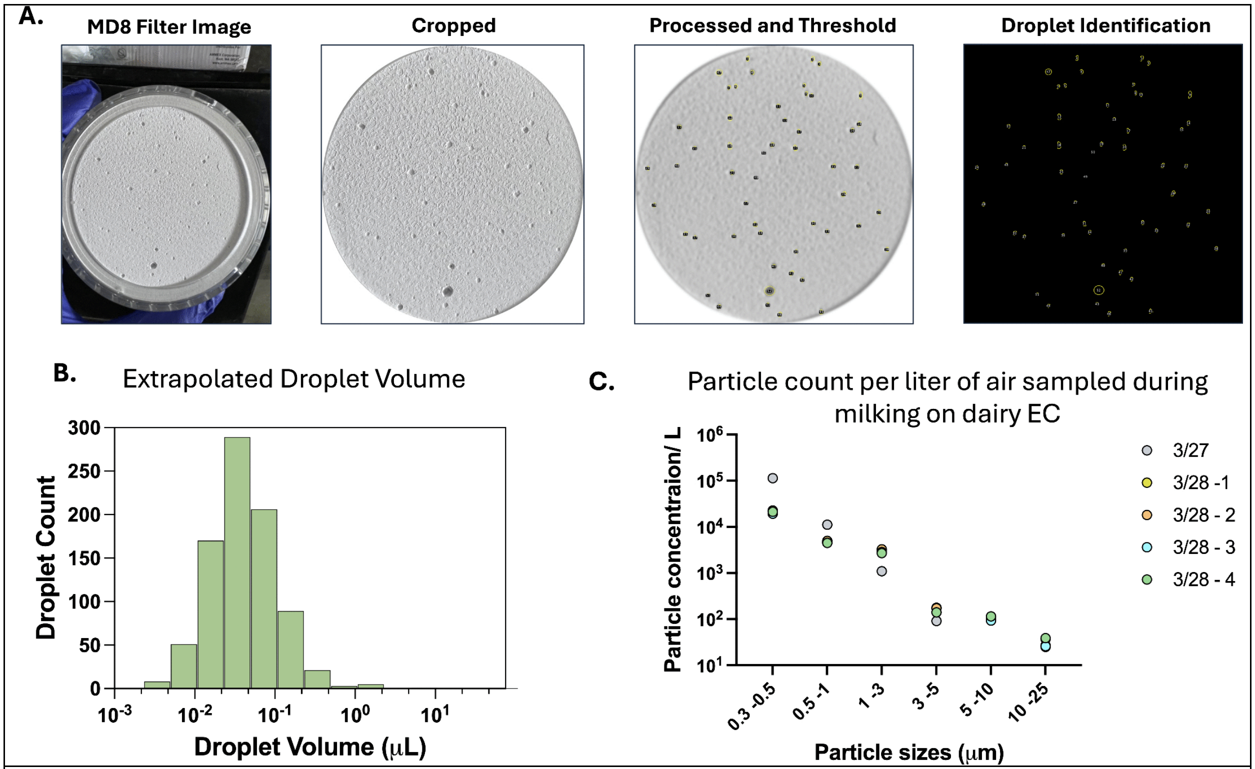


**S1 Fig: Quantification of aerosol and droplet sizes captured in milking parlors.** (A) A representative MD8 gelatin filter with aerosol and droplet marks. Images from milking parlor sampling were collected on multiple days and processed to quantify the aerosol sizes. Processing included cropping of images, blurring, and thresholding of pixel intensities to identify individual spots. (B) Analysis of 14 distinct MD8 filters provided 843 unique spots that were analyzed for the volume of each droplet using an equation to estimate the spread factor (see Materials and Methods). (C) Quantitation of number of aerosol particle counts of different sizes within the milking parlor at farm EC on the indicated sampling dates. Four different measurements were collected on 3/28.

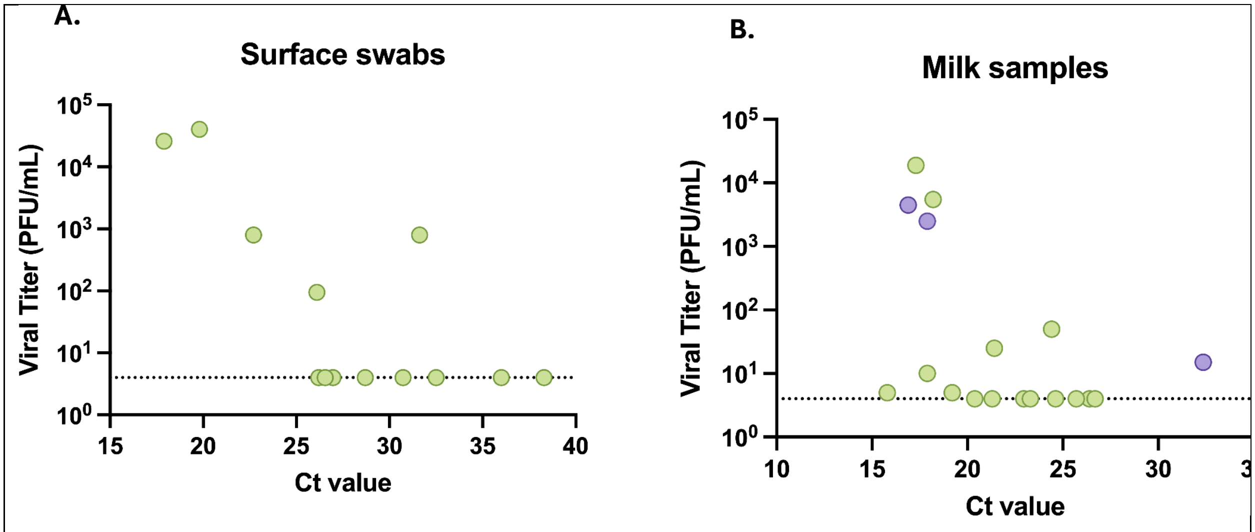


**S2 Fig: Viral loads from milking equipment surface samples and from sick cow milk samples collected prior to initiation of the longitudinal study (prior to 3/18/25).** Infectious virus titers as determined by plaque assay compared to viral RNA levels (Ct values) for (A) swab samples collected from milking equipment or (B) milk collected from sick cows, whose milk would go into buckets and not the bulk tank, on farms EC (green dots) and EG (purple dots).

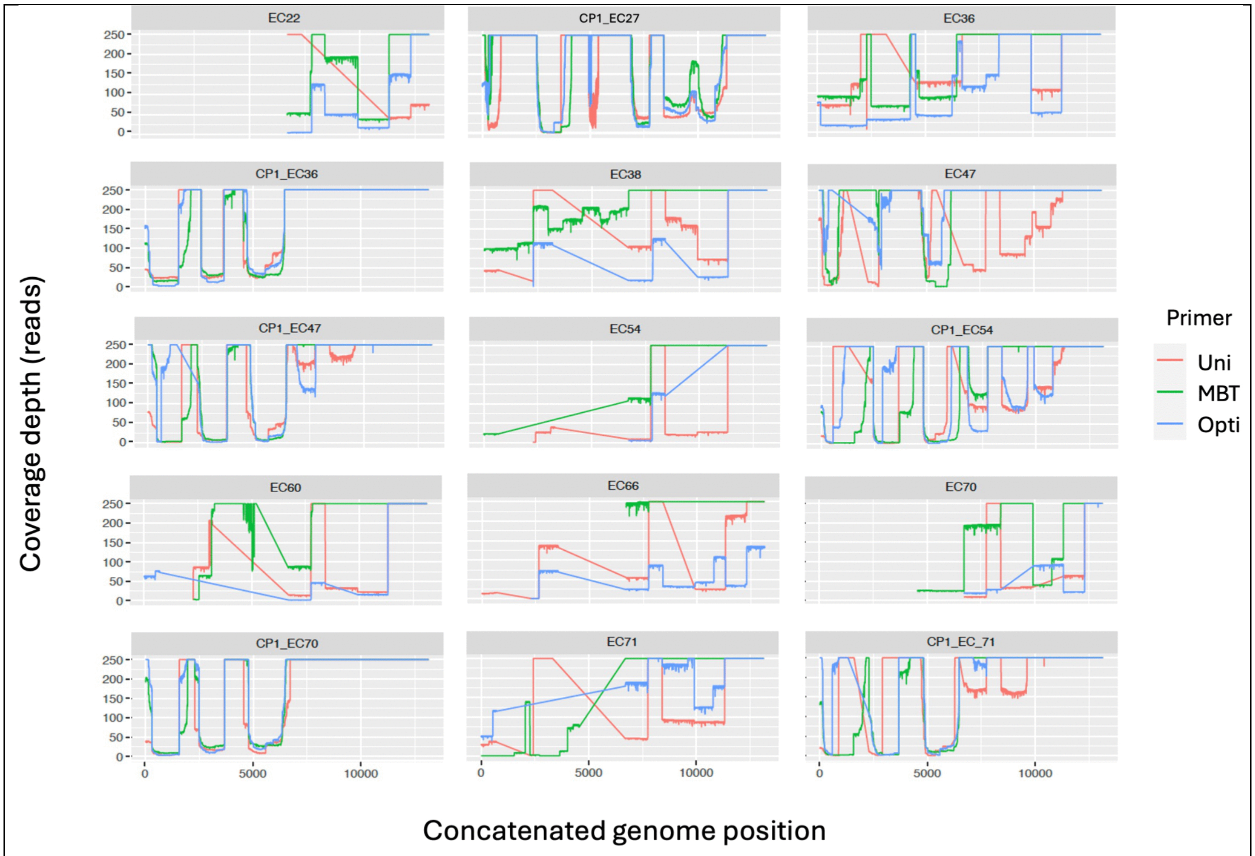


**S3 Fig: Coverage plots for sequencing of environmental samples from dairy farms**. The sequencing read coverage across the concatenated influenza A virus genome for each sample listed in Table 1 is presented in its own plot. Samples sequenced from cell-passaged virus are denoted CP1 for "cell passage 1". The concatenated genome includes PB2 (positions 1-2314), PB1 (positions 2315-4631), PA (positions 4632-6840), HA (positions 6841-8587), NP (positions 8588-10094), NA (positions 10095-11525), M (positions 11526-12527), and NS (positions 12528-13388). Positions with coverage greater than 250 reads were assigned values of 250 on these plots for ease of visualization. Three different primer sets (Uni: Uni12/Inf1, Uni12/Inf3, and Uni13/Inf1[42]; MBT: MBTuni-12, MBTuni-12.4, and MBTuni-13[43]; and Opti: OptiF1, OptiF2, and OptiR[44]) were used to amplify viral genomes prior to sequencing. Variants were only reported if present in the consensus sequence from all primer sets that had coverage at that locus with at least one primer set yielding a coverage depth of at least 24 reads.
