## Supplementary material for "Surveillance on California dairy farms reveals multiple possible sources of H5N1 transmission": Tables S1-S6

**S1 Table. Details of 14 dairy farms that were sampled in California.**

| Dairy Farm | Location (region) | Date of H5 positivity | Approx. Number of Cows in Milk[a] | Primary Housing Pen Style | Milking Parlor Style |
|---|---|---|---|---|---|
| **BB** | Central Valley | 11/23/24 | 800 | Freestall | Parallel |
| **BC** | Central Valley | 10/24/24 | 2900 | Freestall | Rotary and Single Flat |
| **BD** | Central Valley | 11/25/24 | 1000 | Freestall | Parallel |
| **BF** | Central Valley | 11/26/24 | 3000 | Freestall | Rotary |
| **BM** | Central Valley | 11/25/24 | 1400 | Freestall | Herringbone |
| **EA** | Southern CA | 3/3/25 | 300 | Open Lot | Herringbone |
| **EB** | Southern CA | 2/4/25 | 970 | Open Lot | Unknown |
| **EC** | Southern CA | 2/24/25 | 420 | Open Lot | Herringbone |
| **ED** | Southern CA | 12/2024* | 500 | Open Lot | Unknown |
| **EE** | Southern CA | 12/2024* | 1700 | Open Lot | Herringbone |
| **EF** | Southern CA | 12/2024* | 1200 | Open Lot | Herringbone |
| **EG** | Southern CA | 2/26/24 | 500 | Open Lot | Herringbone |
| **FA** | Central Valley | 11/11/24 | 2,000 | Freestall | Herringbone |
| **FB** | Central Valley | 11/19/24 | 3,342 | Freestall | Rotary |

* Approximate date provided

**a** - Cows in milk refers to the number of cows currently lactating and being milked on a dairy farm, not the total number of animals on a farm.

**S2 Table. Details of air sampling during the initial phase on farms in the Central Valley of California in late 2024.**

| Air Sampler | Modification | Sample Location | Sample Descriptor | Farm | Date | Days post BTM+[a] | Positive/ Total |
|---|---|---|---|---|---|---|---|
| AirPrep Cub 210 (200 LPM) | None | Milking Parlor | Cows from multiple pens during milking. Sample duration 60 minutes | BC | 10/29/24 | 5 | 0/4 |
| | | | | BC | 10/30/24 | 6 | 0/3 |
| | | | | BC | 11/4/24 | 11 | 0/2 |
| | | | | BC | 11/7/24 | 14 | 0/1 |
| | | | | BC | 11/11/24 | 18 | 0/2 |
| | | | | BM | 12/4/24 | 9 | 0/2 |
| | | Housing Area[b] | Open-air pen of hospital cows[c] Sample duration 60 minutes | BM | 12/4/24 | 9 | 0/1 |
| Open Face PTFE (5 LPM) | Backpack[d] | Milking Parlor | Following milking process during milking of hospital pen[c] cows. Sample duration 20- 45 minutes | BC | 10/29/24 | 5 | 0/3 |
| | | | | BC | 10/30/24 | 6 | 0/3 |
| | | | | BC | 11/4/24 | 11 | 0/1 |
| | | | | BB | 12/16/24 | 23 | 0/1 |
| | | | Following milking process during milking of cows recovered from clinical H5 signs | BD | 12/18/24 | 23 | 0/1 |
| | | | Following the parlor worker closely during milking of hospital pens of cows | BM | 12/19/24 | 24 | **1/1** |
| | | Housing Area | Sampling of background air in housing pens. Sample duration 20- 45 minutes | BC | 10/29/24 | 5 | 0/1 |
| | | | | BB | 12/16/24 | 23 | 0/1 |
| | | | | BF | 12/18/24 | 22 | 0/1 |
| | Cone[e] | Milking Parlor | Cone directed at milking process, following worker during milking of healthy cows | BB | 12/16/24 | 23 | 0/1 |
| | | | Cone directed at milking process, following worker during milking of cows recovered from H5 clinical signs | BD | 12/18/24 | 23 | 0/1 |
| | | Housing Area | Exhaled breath of 15 - 30 cows in the hospital pen; cone held close to muzzles. Sample duration 5- 15 seconds/cow | BM | 12/16/24 | 21 | 0/1 |
| | | | | BF | 12/18/24 | 22 | 0/1 |
| | | | | BM | 12/19/24 | 24 | 0/2 |
| | | | Exhaled breath of individual cow with 105F fever on 12/16/25 in hospital pen. Sample duration 2 minutes | BM | 12/16/24 | 21 | 0/1 |
| | | | | BM | 12/19/24 | 24 | 0/1 |
| | No Cone | Housing Area | Exhaled breath of individual hospital pen cows. Sample duration 5- 15 minutes | BC | 10/29/24 | 5 | 0/6 |
| | | | | BC | 10/30/24 | 6 | 0/8 |
| | | | | BC | 11/4/24 | 11 | 0/2 |
| MD8 Airport (50 LPM) | Cone | Milking Parlor | Cone directed at milking process, following worker, during milking of a healthy pen of cows. Sample duration 10 minutes | BB | 12/16/24 | 23 | 0/2 |
| | | | Cone directed at milking process, following worker, during milking of the hospital pen. Sample duration 10 minutes | BB | 12/16/24 | 23 | 0/2 |
| | | | Cone directed at milking process, following worker during milking of cows recovered from H5 clinical signs. Sample duration 10 minutes | BD | 12/18/24 | 23 | 0/1 |
| | | | Following the milking process for hospital pens of cows; closely following parlor worker throughout. Sample duration 5- 15 minutes | BM | 12/19/24 | 24 | **3/3** |
| | | Housing Area | Exhaled breath of 15-30 cows, cone held close to muzzles. Sample duration 5- 15 seconds/cow | BB | 12/16/24 | 23 | 0/1 |
| | | | Exhaled breath of 15 - 30 clinically recovered cows; cone held close to muzzles. Sample duration 5- 15 seconds/cow | BD | 12/18/24 | 23 | 0/1 |
| | | | Exhaled breath of 15-30 hospital pen cows; cone held close to muzzles. Sample duration 5- 15 seconds/cow | BF | 12/18/24 | 22 | 0/1 |
| | | | | BM | 12/17/24 | 22 | 0/2 |
| | | | | BM | 12/19/24 | 24 | **2/2** |
| | | | Exhaled breath of individual cow in hospital pen that had an H5+ nasal swab on 12/4/24. Sample duration 1- 2 minutes | BM | 12/16/24 | 21 | 0/2 |
| | | | Exhaled breath of individual cow with severe signs associated with H5, cow couldn't stand. Sample duration 2- 3 minutes | BF | 12/18/24 | 22 | 0/1 |
| | | | Exhaled breath of individual cow in hospital pen that had a 105F fever on 12/16/24. Sample duration 1-2 minutes | BM | 12/19/24 | 24 | 0/1 |

a- Days post BTM+ - Days post first bulk tank milk positive

b - Housing Area refers to primary pens used to house cows; these can be open-air or freestall pens depending on individual dairy.

c - Hospital cows/pen refers to animals identified by farmers, according to internal criteria, that have clinical signs requiring their milk not go to the bulk tank. These animals are grouped into 'hospital pens' separated from healthy animals.

d - Backpack refers to wearing the PTFE filter as shown in Fig 1D; mimicking occupational exposure

e- Cone refers to using a plastic cone adapted to the front of the air sampler as shown in Fig 1D, to prevent aerosol dilution and to protect filters from direct splashes.

**S3 Table. Air sampling details for dairies with 1 or less positive environmental samples, Feb-Apr 2025**

| Sample Location | Sample Type | Sample Source | Sample Descriptor | Farm | Date | Days post BTM+[a] | Positives/ Total |
|---|---|---|---|---|---|---|---|
| Milking Parlor | Milk | Bulk Tank | Collection from bulk tank | EA | 2/26/25 | -5 | 0/1 |
| | | | | EA | 2/27/25 | -4 | **1/1** |
| | | | | EA | 3/4/25 | 1 | **1/1** |
| | | | | EE | 3/2/25 | ~ 90 | 0/1 |
| | | | | FA | 4/1/25 | 141 | 0/1 |
| | | | | FA | 4/2/25 | 142 | 0/1 |
| | | | | FA | 4/3/25 | 143 | 0/1 |
| | | | | FB | 4/1/25 | 133 | 0/1 |
| | | | | FB | 4/2/25 | 134 | 0/1 |
| | | | | FB | 4/3/25 | 135 | 0/1 |
| | | Milk from individual cows, 4 teats | Collected from cows sorted into the hospital pen[b] for signs such as mastitis and potential *E. coli* infection | EF | 3/2/25 | ~90 | 0/3 |
| | | | Collected from cows with abnormal milk consistent with descriptions of milk from H5+ cows such as yellowy color and thick consistency. | FB | 4/2/25 | 134 | 0/2 |
| | | Bulk Sick Cow Milk | Bulk milk from cows whose milk didn't go to bulk tank. No specific clinical signs described | FA | 4/1/25 | 141 | 0/1 |
| | | | Bulk milk from cows whose milk didn't go to bulk tank. Abnormal milk only clinical sign described. | FB | 4/2/25 | 134 | 0/1 |
| | Air | Milking Process | MD8 Airport: cone directed at milking process; following worker. Sample duration 5- 15 minutes | EA | 2/26/25 | -5 | 0/1 |
| | | | | EF | 3/2/25 | ~90 | 0/1 |
| | | | Open Face PTFE worn on backpack following worker during milking. Sample suration 58 minutes | EE | 3/2/25 | ~90 | 0/1 |
| | Surface Swab | Milking Unit Inflations | Swab of the interior of all 4 inflations of one milking unit | EE | 3/2/25 | ~90 | 0/2 |
| Wastewater Stream | Air | Milk Line Cleanout | MD8 Airport held at close range to where wastewater from the milk lines exits the parlor to gravity-flow to the 'flush pump pit' below the manure lagoon. Sample duration 7 minutes | FB | 4/2/25 | 134 | 0/1 |
| | | Sump pump | MD8 Airport: primary sump pump sampled while water flowing. Sample duration 7 minutes | EA | 2/26/25 | -5 | 0/1 |
| | | | MD8 Airport lowered into large, deep sump pump while wastewater flowing. Sample duration 6 minutes | FA | 4/1/25 | 141 | 0/1 |
| | | | | FA | 4/2/25 | 142 | 0/1 |
| | | | Open Face PTFE: primary sump pump sampled while water flowing. Sample duration 12 minutes | EA | 2/27/25 | -4 | 0/1 |
| | | Flush pump pit[c] | MD8 Airport held over area of most agitation of the flush pump pit. Sample duration 5 minutes | FB | 4/1/25 | 133 | 0/2 |
| | | | | FB | 4/2/25 | 134 | 0/1 |
| | | | | FB | 4/3/25 | 135 | 0/1 |
| | | Manure Lagoon | MD8 Airport held, or suspended on a telescoping pole, over inlet for milking parlor wastewater into the lagoon. Sample duration 5- 9 minutes | EA | 2/26/25 | -5 | 0/1 |
| | | | | ED | 3/2/25 | ~90 | 0/1 |
| | | | | EE | 3/2/25 | ~90 | 0/1 |
| | | | | EF | 3/2/25 | ~90 | 0/1 |
| | | | | FA | 4/1/25 | 141 | 0/1 |
| | | Flush of Freestall Pens[d] | MD8 Airport held right in front of where water is pumped up into the freestall pens, vigorous water flow. Sample duration 5- 8 minutes | FB | 4/1/25 | 133 | 0/2 |
| | | | | FB | 4/2/25 | 134 | 0/1 |
| | | | | FB | 4/3/25 | 135 | 0/1 |
| | Wastewater | Milk Line Cleanout | Sample of residual milk and wastewater flushed out of lines as part of the cleaning process post- milking | ED | 3/1/25 | ~90 | 0/1 |
| | | | | EF | 3/2/25 | ~90 | 0/1 |
| | | | | EE | 3/2/25 | ~90 | 0/1 |
| | | | | FB | 4/2/25 | 134 | 0/1 |
| | | Sump pump | 1L from primary pump while water flowing | EA | 2/26/25 | -5 | 0/1 |
| | | | | EA | 2/27/25 | -4 | 0/2 |
| | | | | EA | 3/4/25 | 1 | 0/1 |
| | | | 1L from secondary pump while water flowing | EA | 2/26/25 | -5 | 0/1 |
| | | | | EA | 2/27/25 | -4 | 0/1 |
| | | | | EA | 3/4/25 | 1 | 0/1 |
| | | | 1L sample from deep pump while wastewater flowing. | FA | 4/1/25 | 141 | 0/2 |
| | | | | FA | 4/2/25 | 142 | 0/1 |
| | | | | FA | 4/3/25 | 143 | 0/1 |
| | | Flush pump pit | 1L sample of wastewater, lots of agitation. | FB | 4/1/25 | 133 | 0/1 |
| | | | | FB | 4/2/25 | 134 | 0/1 |
| | | | | FB | 4/3/25 | 135 | 0/1 |
| | | Manure Lagoon | 1L sample from pipe inlet to lagoon | EA | 3/4/25 | 1 | 0/1 |
| | | | | ED | 3/2/25 | ~90 | 0/1 |
| | | | | EE | 3/2/25 | ~90 | 0/1 |
| | | | | EF | 3/2/25 | ~90 | 0/1 |
| | | | 1L sample from lagoon milking parlor wastewater was flowing into | FA | 4/1/25 | 141 | 0/3 |
| | | | | FA | 4/2/25 | 142 | 0/1 |
| | | | | FA | 4/2/25 | 143 | 0/1 |
| | | Field | 1L sample from pump adding a 1:1 mix of wastewater from the manure lagoon to well water onto in-use crop field | FA | 4/1/25 | 141 | **1/1** |
| | | Flush of Freestall Pens | 250mL sample of water collected as it flowed out into the pens. | FB | 4/1/25 | 133 | 0/1 |
| | | | | FB | 4/2/25 | 134 | 0/1 |
| | | | | FB | 4/3/25 | 135 | 0/1 |
| Housing Pens | Air | Exhaled Breath of Row of Cows | MD8 Airport held very close to ~ 12 cows' muzzles as they were headlocked into stanchions. Hospital pen sampled. Sample duration 10- 30 seconds/cow | EF | 3/2/25 | ~90 | 0/1 |

a- Days post BTM+ - Days post first bulk tank milk positive

b - Hospital cows/pen refers to animals identified by farmers, according to internal criteria, that have clinical signs requiring their milk not go to the bulk tank.

c - The flush pump pit is a large basin that was used to mix wastewater from the manure lagoon with fresh water. Water from this pit is used to clean other areas of the dairy, such as freestall pens.

d - The freestall pens are flushed daily with water to clean them out. On the sampled farm this water came from the flush pump pit.

**S4 Table. Sampling details for dairy farm EB during spring 2025.**

| Sample Location | Sample Type | Sample Source | Sample Descriptor | Date | Days post BTM+[a] | Positives/ Total |
|---|---|---|---|---|---|---|
| Wastewater Stream | Air | Sump pump | MD8 Airport (50 LPM) sampling for 7 minutes while pump cycled | 2/26/25 | 22 | 0/1 |
| | | | | 2/27/25 | 23 | 0/1 |
| | | Field | MD8 Airport (50 LPM) sampling over the wastewater outlet in the field for 4- 6 minutes | 2/26/25 | 22 | 0/2 |
| | | | | 3/1/25 | 25 | 0/1 |
| | | | | 3/4/25 | 28 | 0/1 |
| | Wastewater | Sump pump | 1L sample from sump pump | 2/26/25 | 22 | **1/1** |
| | | | | 2/27/25 | 23 | **1/1** |
| | | Field | 1L sample from wastewater inlet to field, usually while pump running | 2/26/25 | 22 | **3/3** |
| | | | | 3/1/25 | 25 | **1/1** |
| | | | | 3/4/25 | 28 | **1/1** |
| | | Manure Lagoon | 1L sample; milkhouse wastewater not flowing here, lagoons only had run-off from housing pens and rainwater | 2/26/25 | 22 | **1/2** |

a- Days post BTM+ - Days post first bulk tank milk positive

**S5 Table. Sampling details for dairy farm EC during spring 2025.**

| Sample Location | Sample Type | Sample Source | Sample Descriptor | Date | Days post BTM+[a] | Positives/ Total |
|---|---|---|---|---|---|---|
| Milking Parlor | Milk | Bulk Tank | Collection from bulk tank | 2/27/25 | 3 | **1/1** |
| | | | | 2/28/25 | 4 | **1/1** |
| | | | | 3/1/25 | 5 | **1/1** |
| | | | | 3/6/25 | 10 | **1/1** |
| | | | | 3/7/25 | 11 | **1/1** |
| | | | | 3/8/25 | 12 | **1/1** |
| | | | | 3/9/25 | 13 | **1/1** |
| | | | | LS: 3/18 - 3/27[b] | 22- 31 | **10/10** |
| | | Sick cow milk - 4 teats combined | Collected from cows with signs such as mastitis or a sudden stop in milk production. | 3/1/25 | 5 | **1/1** |
| | | | | 3/4/25 | 8 | **1/2** |
| | | | | 3/6/25 | 10 | **5/5** |
| | | | | 3/7/25 | 11 | **5/6** |
| | | | | 3/9/25 | 13 | **2/2** |
| | | Dump bucket | Collection from communal bucket used for cows whose milk was discarded. | 3/7/25 | 11 | **1/1** |
| | Air | Milking process, following worker | MD8 Airport (50 LPM) with cone directed at milking process, closely following worker. Sample duration 3- 15 minutes. | 2/28/25 | 4 | **1/4** |
| | | | | 3/1/25 | 5 | **1/1** |
| | | | | 3/4/25 | 8 | **1/1** |
| | | | | 3/6/25 | 10 | **4/5** |
| | | | | 3/7/25 | 11 | **4/4** |
| | | | | 3/8/25 | 12 | **4/4** |
| | | | | 3/9/25 | 13 | **3/4** |
| | | | | LS: 3/18 - 3/27[b] | 22- 31 | **5/14** |
| | | Milking process, following worker | Open face PTFE (5 LPM) worn on backpack during parlor sampling while milking going on. Sampling duration from 3 hours 11 minutes to 4 hours 47 minutes. | 2/28/25 | 4 | 0/1 |
| | | | | 3/6/25 | 10 | **1/1** |
| | | | | LS: 3/18 - 3/27[b] | 22- 31 | 0/4 |
| | Surface Swab | Milking unit Inflations | Swab of the interior of all 4 inflations of one milking unit. | 2/28/25 | 4 | **1/4** |
| | | | | 3/1/25 | 5 | **1/1** |
| | | | | 3/4/25 | 8 | **1/1** |
| | | | | 3/6/25 | 10 | **2/2** |
| | | | | 3/7/25 | 11 | **1/1** |
| | | | | 3/8/25 | 12 | **2/2** |
| | | | | 3/9/25 | 13 | **1/1** |
| | | Milking unit shells | Swab of the exterior of all 4 shells, or teatcups, of a milking unit. | 3/1/25 | 5 | **1/1** |
| | | | | 3/6/25 | 10 | **1/1** |
| | | | | 3/7/25 | 11 | **1/1** |
| | | | | 3/8/25 | 12 | **1/2** |
| | | | | 3/9/25 | 13 | 0/1 |
| Wastewater Stream | Air | Manure Lagoon | MD8 Airport (50 LPM) at close range to point of outlet for wastewater into the manure lagoon. Sampling from 3- 6 minutes. | 2/27/25 | 3 | 0/1 |
| | | | | 3/1/25 | 5 | 0/1 |
| | | | | 3/4/25 | 8 | 0/1 |
| | | | | LS: 3/18 - 3/27[b] | 22- 31 | 0/1 |
| | | Sump pump | MD8 (50 LPM) held over an open sump pump pit as wastewater from the milking parlor line cleaning flowed through. Sampling for 13 minutes. | 3/4/25 | 8 | **1/1** |
| | Wastewater | Milk line cleanout | Sample of bulk milk flushed out of lines as part of the cleaning process post-milking | 3/4/25 | 8 | 0/1 |
| | | | | 3/9/25 | 13 | **1/1**[c] |
| | | | | LS: 3/18 - 3/27[b] | 22- 31 | **7/7** |
| | | Sump pump | 1L sample of wastewater from sump pump pit | 3/4/25 | 8 | **1/1** |
| | | | | LS: 3/18 - 3/27[b] | 22- 31 | **4/7** |
| | | Manure Lagoon | 1L sample sample from right next to wastewater inlet to lagoon | 2/27/25 | 3 | 0/1 |
| | | | | 3/1/25 | 5 | **1/1** |
| | | | | 3/4/25 | 8 | **1/1** |
| | | | | LS: 3/18 - 3/27[b] | 22- 31 | **7/7** |
| Housing Pens | Air | Exhaled Breath of Row of On-Study Cows[d] | MD8 Airport (50 LPM) held very close to cows' muzzles. 10- 30 seconds per cow. | LS: 3/18 - 3/27[b] | 22- 31 | 0/3[e] |

a- Days post BTM+ - Days post first bulk tank milk positive
b - LS, Longitudinal Study. Samples collected between 3/18/25 and 3/27/25 are aggregated.
c - This sample analyzed via qRT-PCR instead of ddPCR, thus it is not shown in Figure 3B.
d - On-study cows refers to the group of 14 cows that were selected for the longevity study wherein daily milk samples from each teat were collected.
e- Not all 14 on-study cows were available for each day sampling conducted

**S6 Table. Sampling details for dairy farm EG during spring 2025.**

| Sample Location | Sample Type | Sample Source | Sample Descriptor | Date | Days post BTM+ [a] | Positives/ Total |
|---|---|---|---|---|---|---|
| Milking Parlor | Milk | Bulk Tank | Collection from bulk tank | 3/6/25 | 8 | **1/1** |
| | | | | 3/7/25 | 9 | **1/1** |
| | | | | 3/8/25 | 10 | **1/1** |
| | | | | 3/9/25 | 11 | **1/1** |
| | | Sick cow milk - 4 teats combined | Collected from cows sorted into the hospital pen that had mastitis at time of sampling. | 3/5/25 | 7 | **1/3** |
| | | Bulk sick cow milk | Milk from sick cows that was diverted from the bulk tank into a smaller tank for a calf raising facility. Clinical signs of cows were decreased appetite and milk production, increased nasal discharge, and increased respiration rates. | 3/6/25 | 8 | **2/2** |
| | Air | Milking process, following worker | MD8 Airport (50 LPM) with cone directed at milking process, closely following worker during milking of the hospital pen cows. Sample duration was 10- 20 minutes. | 3/5/25 | 7 | 0/1 |
| | | | | 3/6/25 | 8 | 0/1 |
| | | | | 3/7/25 | 9 | 0/1 |
| | | | | 3/8/25 | 10 | **1/1** |
| | | | | 3/9/25 | 11 | **1/1** |
| Wastewater Stream | Wastewater | Field | 1L sample from right next to pipe where wastewater from the parlor outlets into the field. | 3/4/25 | 6 | **1/1** |
| Housing Pens | Air | Exhaled breath of row of cows | MD8 Airport (50 LPM) held very close to cows' muzzles as they were headlocked into stanchions. Hospital pen sampled. ~15- 30 cows were sampled for 10- 30 seconds per cow. | 3/7/25 | 9 | 0/2 |
| | | | | 3/8/25 | 10 | 0/1 |
| | | | | 3/9/25 | 11 | **2/3** |

a- Days post BTM+ - Days post first bulk tank milk positive
